# Adult honey bee queens consume pollen and nectar

**DOI:** 10.1101/2024.12.04.626851

**Authors:** Ashley L. St. Clair, Bridget Dwyer, Madeleine Shapiro, Adam G. Dolezal

**Affiliations:** University of Illinois Urbana-Champaign, Urbana, IL 61801, USA

**Keywords:** honey bee, pollinator, pesticide risk assessment, nutrition

## Abstract

Despite the queen’s crucial reproductive role in honey bee colonies, queen diet and feeding behavior remain remarkably enigmatic, with most studies assuming they are solely fed nutritious glandular secretions (i.e., royal jelly) by workers. This colors our understanding of basic honey bee biology and how governmental agencies assess pesticide risk. We hypothesized that adult queens also consume honey and pollen. Through experiments with queenright laboratory microcolonies fed with marked diets, we demonstrate that queens are fed pollen and nectar by workers and can also feed directly. We then measured pollen content in mature, unmanipulated queens sacrificed from 43 conventional field colonies from two distinct geographical regions. Similar to workers, we found pollen in almost all queens guts, though at expectedly lower quantities than in young workers. These findings suggest queens have a more complex, dynamic diet than previously thought, raising new questions about how dietary habits and feeding behaviors influence pesticide risk and other aspects of queen biology.

## Introduction

Nutrition is key to understanding many aspects of animal ecology, development, physiology, behavior, and health. Diet also serves as a major route of agrochemical/pesticide exposure, and regulatory risk assessments model and predict risk by combining exposure, food consumption, and effect data (1). In honey bees, which are also used as a surrogate for other pollinators, worker exposure through contaminated nectar and pollen is well documented (2), informed by studies on their feeding behavior and nutrition (3). While workers perform most colony tasks, each colony contains only one reproductive queen. Substantial research exists on larval queen development, and adult queens can respond rapidly to group-level nutritional changes (4), disease (5), and chemical stress (6). Yet, the diet and feeding behavior of queens remains remarkably mysterious – what do adult queens eat? While queens sometimes consume honey (7, 8), early studies found no evidence of pollen in queen guts (9, 10), leading most research to assume they primarily consume glandular secretions (i.e., royal jelly) fed by workers (11).

This assumption colors our view on many aspects of queen biology and has direct implications on how we predict risk to bees. Because royal jelly is mostly free from pesticide contamination (12, 13), it is assumed that queen bees are insulated from consumption-based exposure. For example, the United States Environmental Protection Agency uses the screening tool BeeREX (1), which incorporates predicted exposure of bees to pesticides and estimates a risk quotient (the ratio of estimated exposure to estimated risk). The BeeREX model can integrate values for consumption of royal jelly, nectar, and pollen to assess risk to different bee developmental stages and castes. For example, worker consumption is modelled based upon maximum consumption rates by age; nurse-age bees are projected to consume 140 mg/day nectar and 9.6 mg/day pollen, and slightly older workers 60 mg/day and 1.7 mg/day, respectively. However, the BeeREX model assumes that queens, whether larval or adult, consume 0 mg/day of nectar and pollen (1), i.e., no exposure through these foods. At a time when beekeepers increasingly report queen losses as a major source of colony loss (14), it is critical to produce risk models which are protective to all castes, and this requires clear assessment of queen diet consumption. Here, we used a mixture of laboratory and field approaches to test the hypothesis that queens do, in fact, consume pollen and nectar, both by feeding themselves and delivery from workers, and thus are subject to calculable risk.

## Results and discussion

Using laboratory microcolony cages, we provisioned solitary queens or queen/worker groups with color-marked sucrose, pollen, both, or unmarked control diets over 7-days. This design enabled non-lethal assessment of diet presence through visual observation through the ventral cuticle over time, as well as confirmation of findings through dissection at the end of the trial (Fig. 1A, B). Solitary queens had direct access to the diets, but in cages with workers, queens were excluded via a screen insert. Overall, we found that queens, whether alone or with attendants, consumed both pollen and nectar, with all marked diet treatments showing significantly higher coloration scores compared to unmarked controls (F_4, 168_=34.11, *p*=<0.0001; Figure 1C). Our solitary queen treatment showed that queens can and do feed directly without worker assistance. Worker-attended queens showed similar coloration scores to our solitary queens, showing that attendant workers are likely feeding queens similar quantities of nectar and pollen.

**Figure 1.**
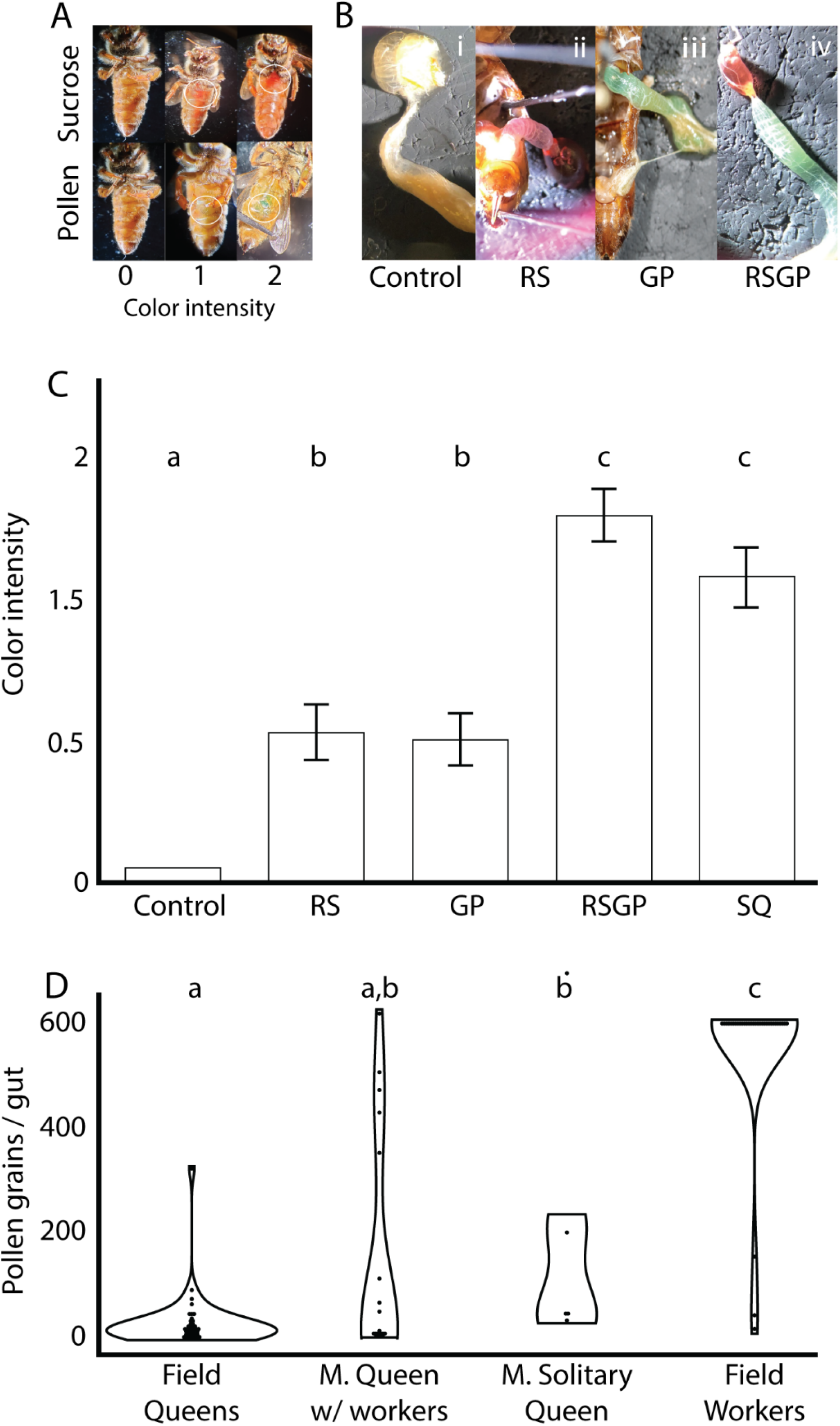
Honey bee queens consume pollen and nectar. **A)** Color intensity scale used to evaluate queen consumption of red-dyed sucrose (top) and green-dyed pollen (bottom) over 7 days; zero scale based on queens fed unmarked diet. Microcolony treatments were control sucrose:pollen (Control), red sucrose:control pollen (RS), control sucrose:green pollen (GP), red sucrose:green pollen (RSGP), and solitary queen (with red sucrose:green pollen (SQ) **B**) Dissection of microcolony queen gut at trial termination validates color scores, showing zero coloration in controls and presence of red sucrose and green pollen in marked diet treatments. **C**) 7-day mean color intensity score (± s.e.) of microcolony treatments; overall treatment resulted in significant differences between groups (mixed model ANOVA, F_4, 168_=34.11, *p*=<0.0001); letters denote pairwise differences between groups (Tukey HSD, *p<*0.05). **D**) Mean (± s.e.) pollen grains in gut of field-collected queens, microcolony (M.) queens with workers, microcolony (M.) queens without workers (i.e., solitary queens), and field-collected young workers; overall group comparison showed significant differences (Kruskal-Wallis test, p<0.001); letters denote pairwise differences between groups (Wilcoxon *post hoc* test, *p<*0.05).

To better understand how these laboratory experiments recapitulate real-world food consumption, we sacrificed queens from 43, healthy, conventionally-managed colonies from apiary systems approximately 120 km apart, in Illinois (n=29) and Louisiana (n=14), USA. For comparison, we also collected putative nurse workers from 33 field colonies and queens from a subset of microcolonies with and without workers (i.e., solitary queens). For all bees, we dissected guts, stained for pollen, and counted grains using microscopy. Contrary to the previous studies finding no pollen in queen guts (9, 10), we found pollen in the vast majority of queens. The proportion of queens containing pollen did not differ between queen types or nurse workers (Pearson’s χ^2^ test (3, N=100)=5.52, p>0.05), supporting our hypothesis that queens commonly consume pollen. When we quantified pollen in the gut, we found significant differences between different queen treatments and workers (Figure 1D, Kruskal-Wallis test, H(3)=60.76; p<0.001). Young workers contained more pollen grains than all other groups (Wilcoxon signed-rank test, p<0.05 for all comparisons). Solitary and worker-attended queens from microcolonies did not differ from each other (Wilcoxon signed-rank test, p>0.05); field-collected queens contained significantly less pollen than solitary microcolony queens (Wilcoxon signed-rank test, p<0.05), but not worker-attended microcolony queens (Wilcoxon signed-rank test, p>0.05). These results show that young workers, who are assumed to consume the most pollen (1), expectedly contain the highest quantities of pollen in their gut; however, while most queens are consuming pollen, different environments may lead to differences in pollen consumption, i.e., variation exists among queens.

## Conclusions

Together, our laboratory and field data support the hypothesis that honey bee queens can and do consume pollen and nectar in addition to their canonical diet of royal jelly. While seemingly simple, these data address a long-standing gap in our understanding and evaluation of adult queen diet and feeding behavior. It also challenges the assumption that queens are not exposed to dietary agrochemical risk. Worldwide agencies, like the European Food and Safety Authority (EFSA) have identified a need to further understand hazard to reproductive queens to accurately assess risk (15), which can only be achieved by better quantifying queen feeding and incorporating these data into predictive models. While thorough comparisons between queens and workers will require more experiments, we can exemplify the importance of identifying these values by roughly contextualizing them within the current BeeREX system. In our queens and workers, guts contained an average of 65.4 and 552 pollen grains, respectively; thus, queens were observed with roughly 11.8% as much pollen as workers. In BeeREX, nurse workers are estimated to consume 9.6 mg pollen/day (1); extrapolating from our data, this would put queen consumption at a little over 1 mg/day, which would place them slightly below ‘comb-building’ workers (1.7mg/day) and above foragers (0.041 mg/day). Thus, we could now improve our risk estimate for queens. Consumption of pollen and nectar sources also influences many aspects of worker biology, changing their response to pathogens, pesticides, and shaping their microbiome (3). If queens are consuming these same foods across space and time, their responses are likely similarly affected, and thus these findings open up many new questions related to queen nutrition. Future work on queen diet may provide many clues to help us understand the foundational biology of bees and help improve pollinator sustainability.

## Methods

We created queenright microcolony cages as in St. Clair et al 2022 (4), with each containing a single, commercially-sourced “Italian” queen, with or without 70 newly-emerged adult workers. Each was provisioned with 30% sucrose-solution and pollen paste (CC Pollen Co.). Food dye (McCormick, USA) was added to some diets to create the following treatments: control sucrose and pollen (Control), red-marked sucrose and control pollen (RS), control sucrose and green pollen (GP), red sucrose and green pollen (RSGP), and solitary queen with both red sucrose and green pollen (SQ); cages were monitored 7 days. Because abdominal cuticle is translucent, coloration of the gut could be assessed daily without harming the queen; color was scored as 0 (no color), 1 (faint color), 2 (intense color) (Fig.1A); a subset were dissected to confirm color intensity metrics matched non-lethal observations (Fig. 1B). A mixed model ANOVA was used to compare color intensity by treatment, day, and their interaction. Treatment and day influenced color intensity, but no interaction was observed, therefore, we compared least squared means with Tukey HSD adjustment by treatment. We also sacrificed 43 mature queens from field colonies (29 from Illinois, USA; 14 from Louisiana, USA, along with putative nurse workers 33 field colonies. Gut contents of queens from the field and microcolony experiments were assessed by dissecting the guts and staining for the presence of pollen with fuchsin stain (4). Due to non-normal data distribution, these data were analyzed via non-parametric Kruskal-Wallis test followed by Wilcoxon signed-rank pairwise tests. For further details on queen gut pollen extraction, and the full dataset, see the supplemental information.

## Supporting information

Dataset for all work

## Acknowledgments

We than Julia Fine for conceptual and technical help with queen microcolonies; Kate Ihle for providing queens and workers from Louisiana apiaries; Nathan Beach for assistance with beekeeping in Illinois. Funding was provided to B. Dwyer by the Camp Family Foundation award at U Illinois, to A. St. Clair from USDA fellowship USDA-WFD 2021-67034-35004, and A. Dolezal through NSF award 2022049.

