## Supplemental methods for "Adult honey bee queens consume pollen and nectar"

**Supplemental Information**

### **Supplemental Methods**

##### ***Bee source***

Worker bees used to supply the laboratory microcolonies were sourced from the University of Illinois (UIUC) Bee Research Facility in Urbana, IL from colonies maintained using standard beekeeping practices including regular *Varroa* mite treatment. To collect day old bees for laboratory cage trials, brood frames were randomly selected from five separate colonies. These frames were stored in individual eclosion boxes in an incubator at 33.5℃ and 50% relative humidity (RH). The emerged bees were brushed from frames daily to ensure each cohort was less than 24 hours old. Bees from all five frames were mixed to randomize colony source effects. From this homogenous mix, we selected a standard weight of bees to fill cages (see below). With remaining bees, an alcohol wash mite check was performed using methods from (1) to ensure that the results of the experiment were not confounded by high mite pressure. Commercially-reared queen bees marketed as Italian (*Apis mellifera ligustica*) were used in all experiments and were acquired from Jackie Park-Burris LLC. in San Pedro, CA.

##### ***Diet preparation***

To better understand whether adult queen honey bees are fed royal jelly exclusively or also fed nectar and pollen directly, we created color-marked sucrose and pollen diets, as well as an uncolored control. To make the sucrose solution, 300 g of granulated white sugar was added to 700 mL of sterilized water (30% sucrose solution w/v); half of this solution (500 mL) remained a control while the other half was treated with 15 drops red colored food dye (McCormick Food Colors & Egg Dye, McCormick & Company, Inc; (2). For all the experiments, bees were fed a mixture of polyfloral pollen from Arizona, USA, purchased from CC Pollen Co. (beepollen.com). To create the pollen diet, 100 g pollen was ground into a fine powder using a coffee grinder, then mixed with 20 g (20% by weight) of 60% (w/v) sucrose solution to create a paste. Half was then used as an uncolored control and the remaining half treated with 20 drops standard blue food coloring. Because of the natural color of the pollen, the resulting dyed pollen color was deep green and referred to as green throughout the paper. Pollen diets were stored at -20°C when not in use.

Using these diets, we set up a combination of queenright microcolonies called queen monitoring cages (QMCs) (see below) in our laboratory, provisioning each with a mix of the control or treated sucrose and pollen diets and with and without the presence of workers.. In total we had five treatment groups; workers present with control sucrose and control pollen (Control), workers present with red sucrose and control pollen (RS), workers present with control sucrose and green pollen (GP), workers present with both red sucrose and green pollen (RSGP), and lastly, a solitary queen (no workers present) with both red sucrose and green pollen (SQ) (Figure 1).

##### ***Laboratory queen monitoring cage set-up***

The QMCs used in these experiments function as queenright microcolonies and mimic colony conditions with the presence of workers, a queen, a feeding chamber which the queen cannot access, and a hexagonal egg laying plate which serves as a space for the queen to lay eggs (3, 4). Each of the five treatments was replicated six times for a total of 30 microcolonies. The trial was run for a total of seven days in July 2021. For treatments where workers were present, cages were filled with 7 g of newly emerged bees (i.e., approximately 70 bees) and all cages received a single Italian queen bee on day zero. Throughout the trial, cages were kept in a dark environmental chamber at 33.5°C and 50% RH.

All QMCs received 2 mL of distilled water, 2 mL of either control or green treated pollen, and 4 mL of either control or red treated sucrose which were added using 2 mL microcentrifuge tubes inserted through holes in the bottom of the cage. When workers were present, the feeding chamber was excluded from the queen egg laying arena by queen excluder material. This allowed for the workers to pass through and care for the queen but denied the queen access to food directly, ensuring the queen was only fed through oral food sharing by workers (i.e., trophallaxis). For the SQ treatment, which has only a queen present, the queen excluder was removed allowing her direct access to the diets. Additionally, we used a pipet to place 100 μL red sucrose within the cells of the egg laying plate to mimic the presence of nectar on a frame. Pollen, water, and sucrose diets were provided *ad libitum* and replenished daily over the seven-day trial.

##### ***Assessment of colored diet consumption in queens***

To determine whether the queen was being fed color treated food or consuming it herself in the absence of workers, we did a visual evaluation of the queen’s abdomen daily. The cuticle of the “Italian” queens is a light gold and when observed from the ventral side, the colored food is distinctly visible in the gut which lies just under the cuticle; this allows observation of colored food consumption without harming the queen. Each day, the queen ventral cuticle was observed through the clear acrylic cage for signs of color in her stomach. The color was rated on an intensity scale from 0-2 (Fig. 1A). A rating of 0 indicated no color could be seen, a rating of 1 indicated faint color was observed, and a rating of 2 indicated the presence of bright color throughout the abdomen. At the end of the seven-day trial, the queens were collected and stored in a -80°C freezer. A subset of 2 queens from each treatment was then later dissected to confirm the presence of control diet, red sucrose, and/or green pollen within the honey stomach and/or gut. Stomachs were photographed using light microscopy. We then used a subset of the remaining queens to dissect the gut open and look for the presence of pollen grains (see below).

##### ***Gut content comparison of queens from field colonies with laboratory microcolonies***

If the queen is fed exclusively a diet of royal jelly, then we would not expect to find pollen within a queen stomach as it is not a component of royal jelly (5). To understand whether queens consume pollen, either on their own or by their attendants, and to validate the results of the laboratory microcolony trial, we collected 29 queens from conventionally managed field colonies at the UIUC bee research facility in Urbana, IL over period spanning August 16^th^, 2022 through August 24^th^, 2023 (8/16/2022 8 queens; 9/16/2022 2 queens; 10/1/2022 2 queens; 5/3/2023 6 queens; and 8/24/2023 11 queens). Additionally, we collected 14 queens from the research apiary of the USDA-ARS Bee Breeding and Genetics laboratory in Baton Rouge, Louisiana, USA on August 2^nd^, 2023. We then compared the gut contents of field collected queens to those reserved from the solitary queen and worker attended queen QMC microcolony trials. Because we did not have high replication of the solitary queens remaining from the colored diet consumption trial (n=4 queens), we ran a second trial, following the same methods as above, of solitary queens in microcolonies starting on July 25, 2023, and replicated this 15 times for a total of 19 replicate solitary queens.

***Measurement of pollen contents via microscopy***

To evaluate pollen in the bee stomach, we first dissected out the honey stomach and gut into a 1.5 mL microcentrifuge tube and added 1.5 mL of 70% ethanol. The gut was then homogenized using a sterile dissecting pin and centrifuged at 4000x rpm for 5 minutes. The ethanol was decanted, leaving a pellet of pollen and broken gut tissue. Using a clean toothpick, the gut pellet was removed and rubbed onto a drop of fusion based gelatin on a microscope slide. The remaining tissue was placed back into the microcentrifuge tube, re-diluted with 1.5 mL of 70% ethanol, and stored at 4°C. For each bee, two replicate stains were made. To quantify pollen, each stain was observed under light microscopy and pollen grains were counted, up to 300 (6). The total of both replicates was summed for a maximum of 600 possible grains counted.

##### ***Statistical analysis***

To assess abdominal color in queens, which was used as a proxy to indicate the level at which queens consumed experimental diet, we created a mixed model analysis of variance in JMP Pro v18.0.1. Queen abdomen color ranking was the response variable with treatment, trial day, and their interaction as fixed effects. There was an overall significant effect in the model (F_34,168_=5.47, p=<.0001) with treatment (F=34.11, p=<.0001) and trail day (F=4.25, p=0.0005) varying significantly, but no interaction was observed between treatment and trial day (F=0.92, p=0.57). Since no interaction was observed, a *post hoc* analysis of least squared means with Tukey HSD adjustment was conducted at the treatment level to compare the overall color intensity of each treatment diet (Fig. 1C).

To compare whether workers and different queen types show different proportions of pollen presence we compared presence/absence of pollen in each type via Pearson’s χ^2^ test. To quantify amount of pollen within queen guts across treatments (i.e., field, cage queen with workers, and solitary queens), we summed the total pollen grains across the two replicates per queen to create a total value. Because these data did not fit the assumptions of homogeneity of variance or normality, they were analyzed via nonparametric statistical methods. First, we tested whether there were differences in field-collected queens from Louisiana and Illinois via Wilcoxon method, finding no differences (Z= -0.38; p=0.71). We then ran Kruskal-Wallis ANOVA on the complete dataset, followed by post hoc pairwise comparisons using the Wilcoxon method. All analyses on gut contents were performed in JMP Pro v.16.
